# Library size confounds biology in spatial transcriptomics data

**DOI:** 10.1101/2023.03.15.532733

**Authors:** Dharmesh D. Bhuva, Chin Wee Tan, Claire Marceaux, Jinjin Chen, Malvika Kharbanda, Xinyi Jin, Ning Liu, Kristen Feher, Givanna Putri, Marie-Liesse Asselin-Labat, Belinda Phipson, Melissa J. Davis

## Abstract

Spatial molecular technologies have revolutionised the study of disease microenvironments by providing spatial context to tissue heterogeneity. Recent spatial technologies are increasing the throughput and spatial resolution of measurements, resulting in larger datasets. The added spatial dimension and volume of measurements poses an analytics challenge that has, in the short-term, been addressed by adopting methods designed for the analysis of single-cell RNA-seq data. Though these methods work well in some cases, not all necessarily translate appropriately to spatial technologies. A common assumption is that total sequencing depth, also known as library size, represents technical variation in single-cell RNA-seq technologies, and this is often normalised out during analysis. Through analysis of several different spatial datasets, we noted that this assumption does not necessarily hold in spatial molecular data. To formally assess this, we explore the relationship between library size and independently annotated spatial regions, across 23 samples from 4 different spatial technologies with varying throughput and spatial resolution. We found that library size confounded biology across all technologies, regardless of the tissue being investigated. Statistical modelling of binned total transcripts shows that tissue region is strongly associated with library size across all technologies, even after accounting for cell density of the bins. Through a benchmarking experiment, we show that normalising out library size leads to sub-optimal spatial domain identification using common graph-based clustering algorithms. On average, better clustering was achieved when library size effects were not normalised out explicitly, especially with data from the newer sub-cellular localised technologies. Taking these results into consideration, we recommend that spatial data should not be specifically corrected for library size prior to analysis unless strongly motivated. We also emphasise that spatial data are different to single-cell RNA-seq and care should be taken when adopting algorithms designed for single cell data.

## Introduction

After being crowned method of the year 2020 ^1^, spatial molecular technologies have advanced drastically with new platforms boasting greater coverage of transcripts and increased spatial resolution of measurements ^2-5^. Resolutions from these technologies span from 100s of cells (e.g., GeoMx) to sub-cellular (e.g., CosMx, Xenium and STOmics); while transcript and protein coverage range from 100s of molecules (e.g., CosMx and Xenium) to genome-wide measurements (e.g., GeoMx, Visium and STOmics). These approaches detect transcripts by either sequencing or imaging, with the latter providing the highest spatial resolution. The ability to resolve high-throughput molecular measurements in space has enabled the study of diseases in their resident tissue microenvironment, thus, providing a more comprehensive view of disease systems ^6^.

The added spatial information coupled with the scale of the data poses a significant bioinformatics challenge. Since it is difficult to conceptualise analysis of individual molecular measurements at sub-cellular spatial resolution, a popular approach has been to abstract the measurements at the cellular level ^3,4^. This is done by segmenting cellular boundaries and accumulating individual datapoints within these cellular bins ^7^. This approach enables the >1300 tools developed for the analysis of single-cell RNA sequencing (scRNA-seq) data to be applied to spatial molecular data ^8^. While applying scRNA-seq tools to spatial molecular data often works well as a first pass ^3,4^, it remains underpowered since these methods disregard spatial information. Dedicated methods that incorporate spatial information are now being developed for analysis tasks such as the identification of spatially variable features ^9-11^, spatially constrained clustering ^12-14^, and cell type annotation ^15,16^. However, these methods are still built on the foundations of cell-based analysis and therefore propagate some of the assumptions inherent to single-cell data. One such assumption is that differences in the total number of transcripts detected/sequenced per cell represents technical variation that should be normalised out prior to downstream analysis. In sequencing-based transcriptomics, this is often referred to as the library size. For imaging-based spatial molecular technologies, it is more appropriate to refer to these as the total detections per cell.

The idea of normalisation for library sizes originated from bulk RNA sequencing where samples were sequenced at varying depths thus the effect of sequencing depth needed to be corrected to enable cross-sample comparison of gene counts ^17^. The simplest method of accounting for library size in RNA-seq data is to divide each count by the total sequencing depth for that sample, and multiply by a scalar, such as a million, to obtain counts per million (CPM), and this has been adopted by the single cell field ^18^. However, sometimes this adjustment does not mitigate the effect of total sequencing depth in single cell experiments and new methods such as regularised negative binomial regression (sctransform)^19^ and the deconvolution of pooled size factors (scran) ^20^ have been proposed to effectively reduce the impact of library size differences. These methods specifically account for the sparsity inherent to single-cell sequencing data. Their application to such data is warranted as each cell is the unit of measurement in these data.

The unit of measurement in sub-cellular spatial molecular technologies is either a transcript detection (e.g., Xenium, CosMx, and FISH-based assays) or a sub-cellular spot (e.g., STOmics) therefore normalisation at the cellular level is not as naturally motivated compared to bulk or scRNA-seq. Although cellular binning is not performed in Visium data, like other spatial molecular technologies, the proximity of spots/cells to neighbouring spots/cells implies spatial autocorrelation resulting from biological dependence when spots/cells originate from the same tissue region. This spatial autocorrelation has not been previously investigated in the context of normalisation for spatial molecular data and is not accounted for in single-cell normalisation methods even though these methods are routinely applied to spatial data from both imaging-based ^21^ and sequencing-based technologies ^2^.

Here, we analyse spatial transcriptomic datasets from four different technologies and three different tissues to show that library size or total detections per cell is not simply a technical artefact that should be corrected for when analysing spatial datasets. Across all four technologies, we show through statistical modelling that library sizes or total detections per cell significantly differ across tissue structures, thus representing real biology rather than technical variation. Similar observations have been made in scRNA-seq data however, this is the first time it has been rigorously tested in spatial molecular data ^22^. We also show that on average, normalising this effect out will negatively impact spatial domain identification. Our recommendation when analysing spatial data is to carefully consider when to normalise library sizes or total detections per cell. For instance, library size normalisation should not be performed prior to spatial domain identification but could be considered for other downstream analytical tasks such as cross-sample comparisons.

While we show that no normalisation outperforms sctransform for clustering tasks, there is clearly a need for new normalisation methods that account for the unique properties of spatial data, such as differences in capture efficiency across the tissue. Here we have specifically evaluated the effect of library size normalisation on clustering, however this could impact the performance of other downstream analysis as well. Overall, we recommend that care is needed when adopting single cell methods to analyse spatial data, as the assumptions of these methods may be violated when applied to spatial data.

## Results

### Library size or total detections per cell captures real biology in spatial transcriptomics datasets

In some single cell datasets with subtle biological signals, library size is often the largest source of variability and can lead to the identification of clusters that capture library size differences, not biology. To assess this in spatial data, we analysed 23 biological samples from 4 different spatial technologies encompassing both imaging- and sequencing-based spatial technologies that span sub-cellular and region-level spatial resolutions. These data are described in Table 1. We began by exploring total detections across space by binning transcript detections from Xenium, STOmics and CosMx into a hexagonal tessellation and visualising the density across bins/spots (Figures 1a-d, Additional File 1: Supplementary Figure 1). These bins were large enough to contain 10s of cells and 1000s of transcripts. To assess library size associations with tissue regions, we independently annotated regions in the Xenium, STOmics and CosMx datasets using immunofluorescence images (see Methods). This allowed us to annotate 149-155 brain regions in the Xenium mouse brain dataset, 118 regions in the STOmics mouse brain dataset and 4 regions in the CosMx non-small cell lung cancer (NSCLC) dataset to enable a comparison across tissue regions (Figures 1e-h, Additional File 1: Supplementary Figure 2). Mouse brain data were annotated using the Allen Brain Atlas ^23^ while the NSCLC data were annotated using QuPath ^24^ to segment regions based on markers.

**Table 1:** Spatial transcriptomics datasets used to study library size effects.

| Technology | Technology type | # samples | # genes | Total counts/ detections | Organism | Tissue | Source |
| --- | --- | --- | --- | --- | --- | --- | --- |
| 10x Visium | Sequencing-based | 12 | 33,538 | 9-22M | Homo sapiens | Dorsolateral prefrontal cortex | <sup>25,26</sup> |
| 10x Xenium | Imaging-based | 3 | 248 | 58-62M | Mus musculus | Brain | <sup>27</sup> |
| NanoString CosMx | Imaging-based | 7 | 960 | 25-40M | Homo sapiens | Non-small-cell lung cancer | <sup>3</sup> |
| BGI STOmics | Sequencing-based | 1 | 26,177 | 134M | Mus musculus | Brain | <sup>2</sup> |

**Figure 1:**
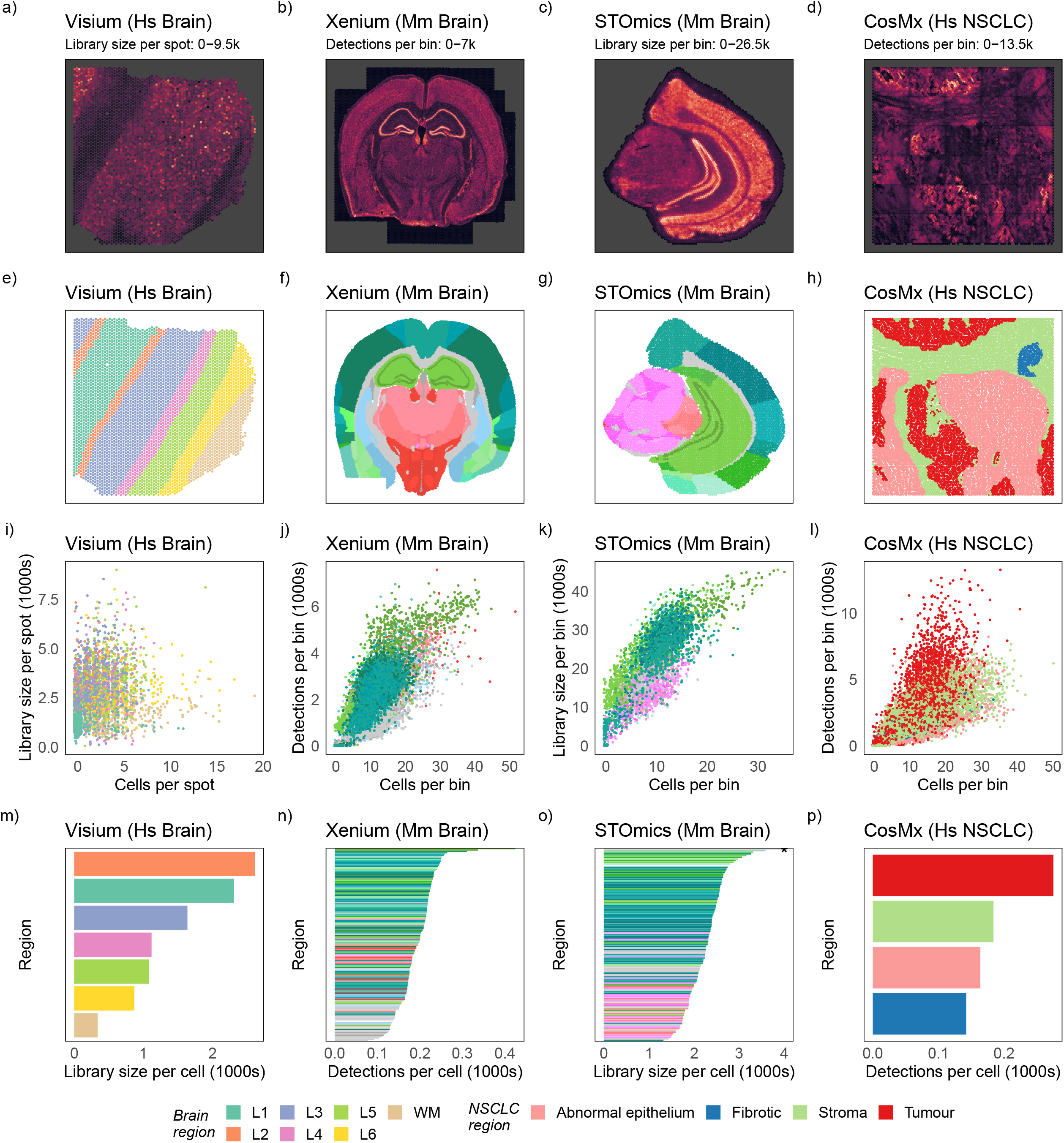
Detection density and total detections/library sizes are associated with biology consistently across different spatial molecular technologies, organs and species. a-d) Detection density per bin/spot plot for Visium dorsolateral prefrontal cortex (DLPFC), Xenium mouse brain, STOmics mouse brain and CosMx non-small cell lung cancer (NSCLC), reveal tissue structure. e-h) Regions annotated for each bin/spot using the Allen Brain Atlas for the mouse brain and manual annotation based on immunofluorescence markers of CosMx NSCLC. i-l) Number of cells plot against the total detections/library sizes per bin/spot, coloured by the tissue region, showing the region-specific relationship between cells and detections/counts. m-p) Average detections/library sizes per cell for each region, computed as the sum of detections divided by the number of cells for each region, showing that related regions exhibit similar total detections/library sizes per cell.

Tissue structure was apparent across the brain and cancer datasets when visualising the total detections/library sizes. With the Visium brain dataset (Figure 1a), we could clearly identify the layering of the cortex (Figure 1e) while with the Xenium and STOmics mouse brain datasets (Figures 1b-c), we could visually identify the cortex (darker greens in Figures 1f-g), white matter (pinks in Figures 1f-g) and hippocampus (brighter greens in Figures 1f-g). Due to the higher spatial resolution of these datasets, we could also identify substructures of the mouse brain such as the dentate gyrus (beak like structure) that had the highest total detections/library sizes. There were clear structures with a large detection count in the NSCLC dataset as well (Figure 1d) with tumour regions having the highest total detections (Figure 1h).

Our binning strategy allowed us to investigate total detections/library size without delving into cell boundary detection which is still an active area of research. However, this meant that each bin contained multiple cells, therefore we had to relate the total detections/library sizes back to the number of cells. As expected, the library size linearly increased with the number of cells regardless of the technology, although this relationship was not as strong for the Visium data (Figures 1i-l, Additional File 1: Supplementary Figure 3). However, we can also see clustering of points by region, particularly for the Xenium and STOmics datasets, indicating that cell density is not the only contributing factor to library size. To demonstrate this effect more clearly, we estimated the total detections/library sizes per cell for each region by dividing the total detections/counts in the region by the total number of cells. Figures 1m-p show each region sorted by these averages across the 4 technologies. We see a clear region-specific effect in each dataset. For the Xenium and STOmics mouse brain datasets, similar brain sub-structures cluster together indicating that the average total detections/library size per cell is similar in the higher-order structures (Figures 1n-o, Additional File 1: Supplementary Figure 4). We also see that tumour regions tend to have higher total detections per cell. This is unsurprising as tumour cells are expected to be transcriptionally more active than other cell types ^22,28^.

Next, we wanted to assess the relationship between regions, the number of cells and total detections/library sizes in a more statistically rigorous manner. To do so, we treated all transcript detections, regardless of the gene, as a spatial point process that is a realisation of an underlying intensity function. This was done by fitting a Poisson model to the total detections/library sizes per bin with the following covariates: cell density, tissue region, and other technology-specific variables such as the field-of-view (CosMx) and the number of DNA nanoball spots (STOmics). The model was fitted to binned data, where the bins are quadrats defined by a hexagonal tessellation. The interaction between all covariates were included in the model. Performing a Type II analysis of variance (ANOVA) ^29^ on the covariates of each model, we found that the number of cells per bin explained the largest variance in library sizes followed by the tissue region (tissue region p-values < 2×10^−308^, Table 2, Supplementary Table 1), across all technologies except for STOmics. In STOmics, the number of DNA nanoball spots was the strongest predictor, however, this number is dependent on the number of cells since nanoball spots not overlapping cells contain no measurements and therefore are not included in the analysis. Collectively, these results show that even after accounting for the number of cells in each bin, there is a significant relationship between spatially defined regions and total detections/library sizes. This effect appears to be technology, species, and organ agnostic, and is present across both healthy and disease systems.

**Table 2:** Results of Type II ANOVA tests on regression models of library size/total detections. (Df – degrees of freedom, Pr(>F) – p-value, Sum Sq – sum of squares)

| Sample | Sum Sq | Df | F value | Pr(>F) | Covariate | Platform |
| --- | --- | --- | --- | --- | --- | --- |
| Human_DLPFC_1 | 34500.725 | 1 | 100.683 | $1.98 \times 10^{-23}$ | NCell | Visium |
| Human_DLPFC_1 | 117179397 | 7 | 48851.871 | $< 2 \times 10^{-308}$ | Region | Visium |
| Human_DLPFC_1 | 8845.691 | 6 | 4.3024 | 0.00024808 | NCell:Region | Visium |
| mBrain_ff_rep1 | 4898911.5 | 1 | 26560.22 | $< 2 \times 10^{-308}$ | NCell | Xenium |
| mBrain_ff_rep1 | 408444642 | 140 | 15817.478 | $< 2 \times 10^{-308}$ | Region | Xenium |
| mBrain_ff_rep1 | 836131.428 | 138 | 32.849 | $< 2 \times 10^{-308}$ | NCell:Region | Xenium |
| STOmics Brain | 5858.959 | 1 | 136.689 | $2.92 \times 10^{-31}$ | NCell | STOmics |
| STOmics Brain | 64978830.5 | 118 | 12847.026 | $< 2 \times 10^{-308}$ | Region | STOmics |
| STOmics Brain | 4230921.64 | 1 | 98706.943 | $< 2 \times 10^{-308}$ | NSpots | STOmics |
| STOmics Brain | 38476.177 | 108 | 8.312 | $5.20 \times 10^{-115}$ | NCell:Region | STOmics |
| STOmics Brain | 62629.932 | 1 | 1461.150 | $3.09 \times 10^{-288}$ | NCell:NSpots | STOmics |
| STOmics Brain | 72415.583 | 109 | 15.500 | $1.57 \times 10^{-245}$ | Region:NSpots | STOmics |
| STOmics Brain | 58914.306 | 104 | 13.216 | $9.20 \times 10^{-197}$ | NCell:Region:NSpots | STOmics |
| Lung5_Rep3 | 4346427.84 | 1 | 10273.966 | $< 2 \times 10^{-308}$ | NCell | CosMx |
| Lung5_Rep3 | 2601294.53 | 4 | 1537.217 | $< 2 \times 10^{-308}$ | Region | CosMx |
| Lung5_Rep3 | 1712746.52 | 29 | 139.605 | $< 2 \times 10^{-308}$ | fov | CosMx |
| Lung5_Rep3 | 24695.567 | 3 | 19.458 | $1.40 \times 10^{-12}$ | NCell:Region | CosMx |
| Lung5_Rep3 | 150009.021 | 29 | 12.227 | $1.03 \times 10^{-56}$ | NCell:fov | CosMx |
| Lung5_Rep3 | 197636.265 | 41 | 11.394 | $5.18 \times 10^{-72}$ | Region:fov | CosMx |
| Lung5_Rep3 | 67589.498 | 40 | 3.9941 | $4.19 \times 10^{-16}$ | NCell:Region:fov | CosMx |

### Normalising out total detections/library sizes reduces clustering performance

Predicated on the region-specific total detections/library size effect, we could infer that normalising out total detections/library sizes would result in loss of information when attempting to identify spatial domains using clustering. This task is commonly performed on Visium data using a standard single-cell clustering pipeline ^30^. This workflow involves normalising out library sizes using sctransform/scran, identifying highly variable genes, performing dimension reduction using principal components analysis, using the top principal components to build a shared nearest neighbour graph, and finally running community detection on these graphs to identify spatial domains.

We wanted to evaluate the impact of normalisation on this workflow without biases in parameter choice. Data normalisation using different methods may mean a different set of parameters work best for each normalisation. To remove any parameter-specific effects, we set up a benchmark that explores a large parameter space and tests all combinations of parameters for each normalisation strategy across 23 samples spanning all four technologies (Figure 2a). In total we tested 14076 different combinations. For each combination of sample and normalisation strategy, we computed the median and maximum Adjusted Rand Index (ARI) representing the average- and best-case scenarios for clustering respectively. Figure 2b shows these values when data were unnormalised, normalised with scran ^20^, or normalised with sctransform ^19^. We see that the median ARI across most samples is higher when no normalisation or scran normalisation is performed than when library size effects are explicity removed using sctransform. This indicates that on average, we are likely to encounter a better clustering without normalisation or when normalising with scran (Figure 2b). If parameters are tuned well in the workflow, scran normalisation can result in better clustering, primarily for the Visium samples. These results again highlight that total detections/library sizes themselves contain region-specific information. Improved clustering on scran normalised data compared to sctransform normalised data could be an indicator of some residual library size effect following scran normalisation since scran does not explicitly normalise out library size effects. We note that this comparison was performed without using spatial localisation information and it would be interesting to see the impact of normalisation on methods dedicated to spatial molecular technologies. Of note is that in general the ARI is not particularly high (max ARI = 0.6), indicating a need for improved analysis methods for spatial data.

**Figure 2:**
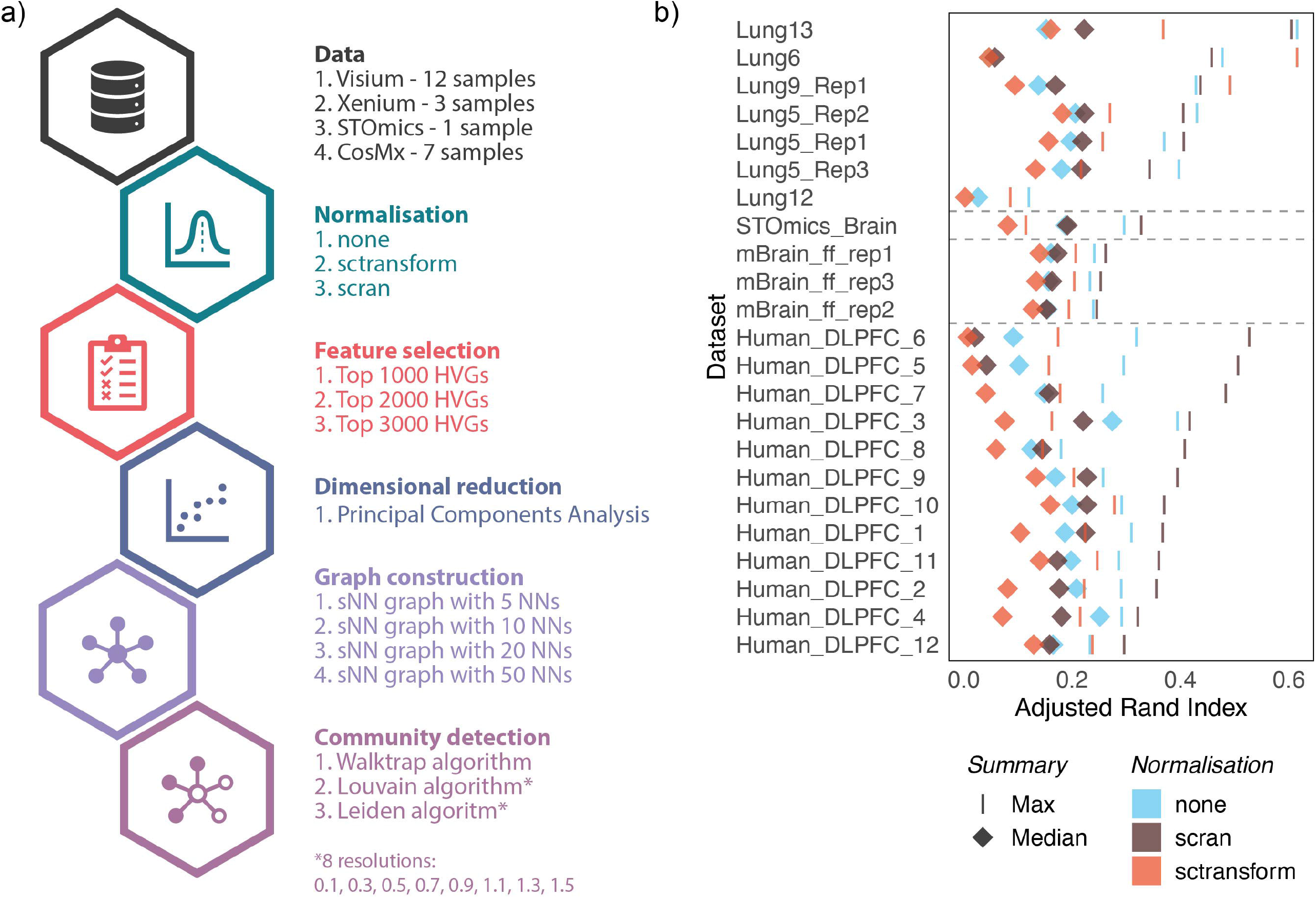
Normalisation of total detections/library sizes results in poorer spatial domain identification using clustering approaches. a) Schematic of the benchmark performed on 12 Visium dorsolateral prefrontal cortex (DLPFC) samples showing the parameter space explored when using a single-cell clustering pipeline to identify spatial domains. b) The median and maximum Adjusted Rand Index (ARI) obtained when no normalisation is performed and when scran and sctransform normalisation are applied. On average, as indicated by the median ARI, no normalisation results in best performance, however, when finely tuned, scran normalisation can produce a better clustering. Specifically, unnormalised data from sub-cellular localised technologies results in better or similar clustering to normalised data.

## Conclusion

Total detections/library sizes are associated with biology in spatial molecular technologies and can be valuable in identifying spatial domains in tissue. They should not be prematurely normalised out as they can enhance various analyses if used carefully. We recommend carefully selecting when to normalise library sizes in spatial molecular data. We also emphasise caution when transferring ideas and tools from single-cell analysis into spatial molecular data as the assumptions of these methods may not be valid for spatial data.

## Methods

### Hexagonal tessellation of sub-cellular localised data

We computed a hexagonal tessellation such that there were 100 hexagons along each axis. Since the area profiled in the Xenium dataset was larger, the tessellation of this dataset contained 200 hexagons along each axis. This was preferred over a standard square grid as a hexagonal tessellation is less prone to edge effects ^31^. Total detections/counts as well as the total number of cells were computed in each bin.

### Poisson model of binned counts

Points in space represent a Poisson point process therefore binning points will result in Poisson distributed count data. We model binned counts as a linear combination of the number of cells, the region types, and any technology specific technical covariates such as the number of DNA nanoball beads (BGI STOmics) and the field of view (NanoString CosMx). Generalised linear models with a log link function are used to perform the fit. All possible interactions between covariates were included in the models.

### Annotating brain datasets using the Allen Brain Atlas

Mouse brain data from the Xenium and STOmics technologies were annotated by registering our DAPI stained images to the common coordinates framework v3 (CCFv3) of the Allen Brain Atlas ^23^ using the Aligning Big Brains & Atlases (ABBA) plugin (v0.3.7) in Fiji (v1.53t) ^32^. The resultant hierarchical annotation was compressed such that the deepest layer of non-missing annotation was used to annotate each detection/DNA nanoball spot. Non-small cell lung cancer (NSCLC) data were annotated manually with QuPath (v0.3.2) ^24^ using the accompanying PanCK, CD3, CD45 and DAPI stained images. Hexagonal bins were then allocated to regions based on the predominant annotation of data points in the bin.

