## Additional File 1 for "Library size confounds biology in spatial transcriptomics data"

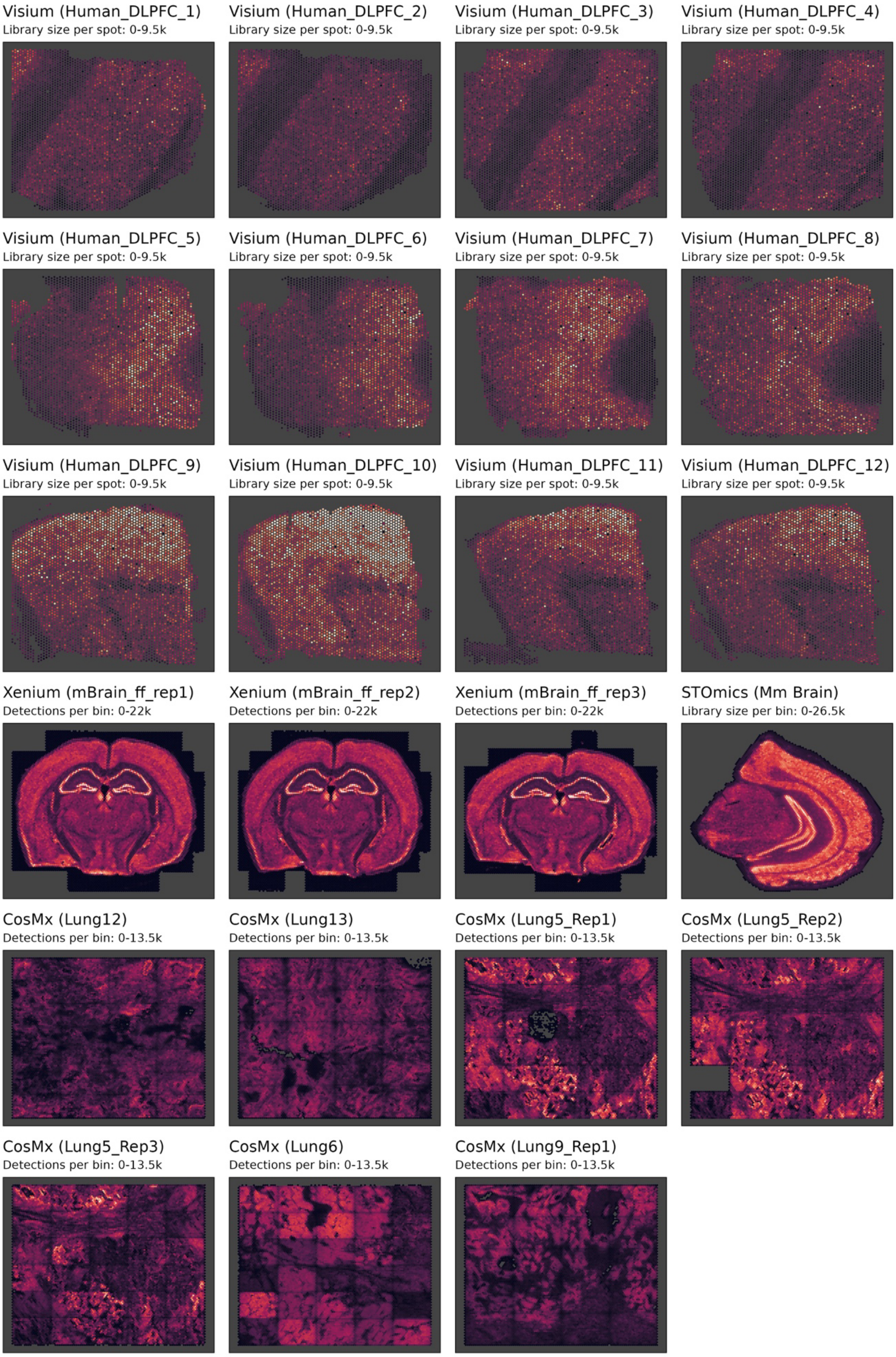


***Supplementary Figure 1:*** *Detection density per bin/spot plot for Visium dorsolateral prefrontal cortex (DLPFC), Xenium mouse brain, STOmics mouse brain and CosMx non-small cell lung cancer (NSCLC), reveal tissue structure.*

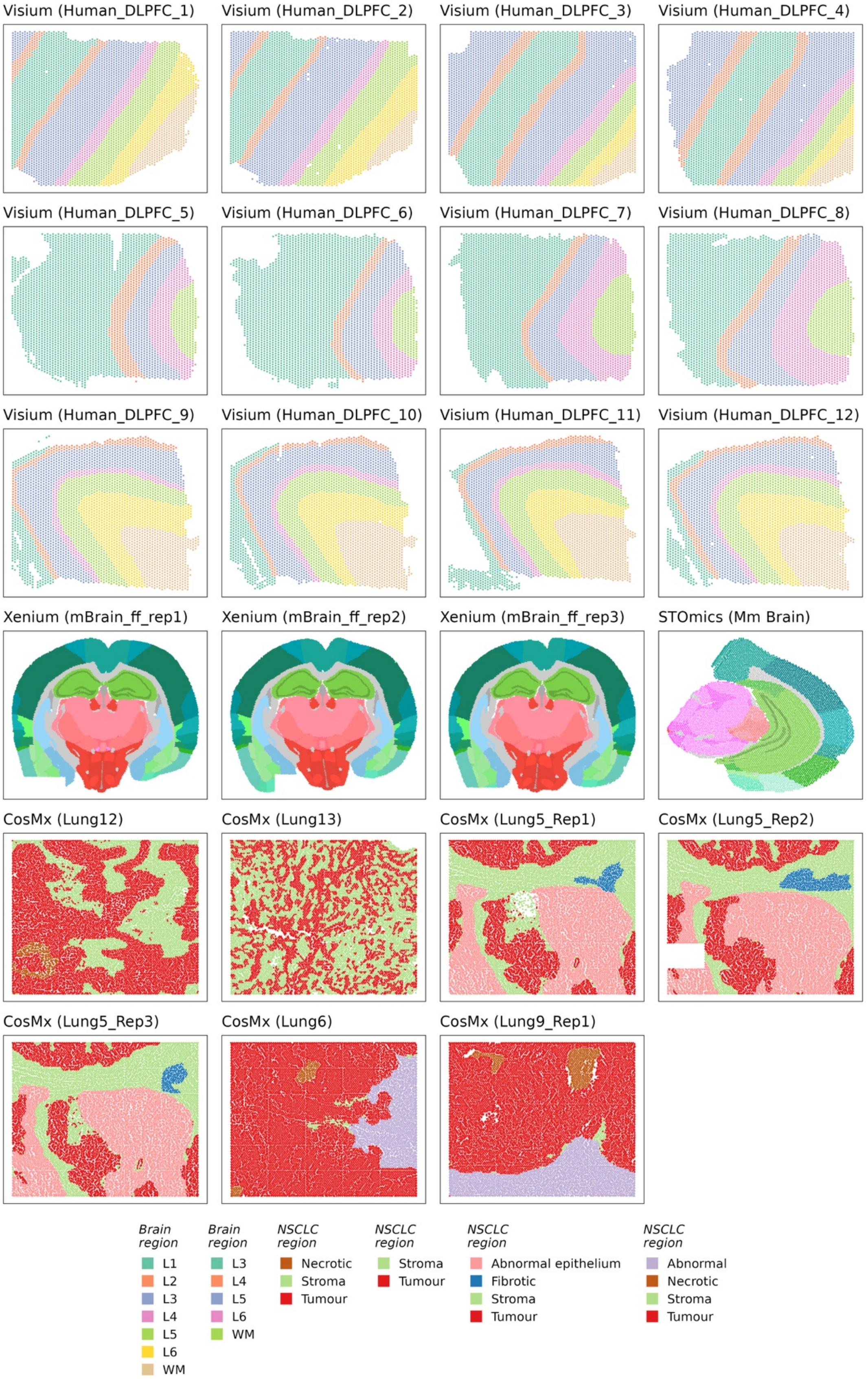


***Supplementary Figure 2:*** *Regions annotated for each bin/spot using the Allen Brain Atlas for the mouse brain and manual annotation based on immunofluorescence markers of CosMx NSCLC.*

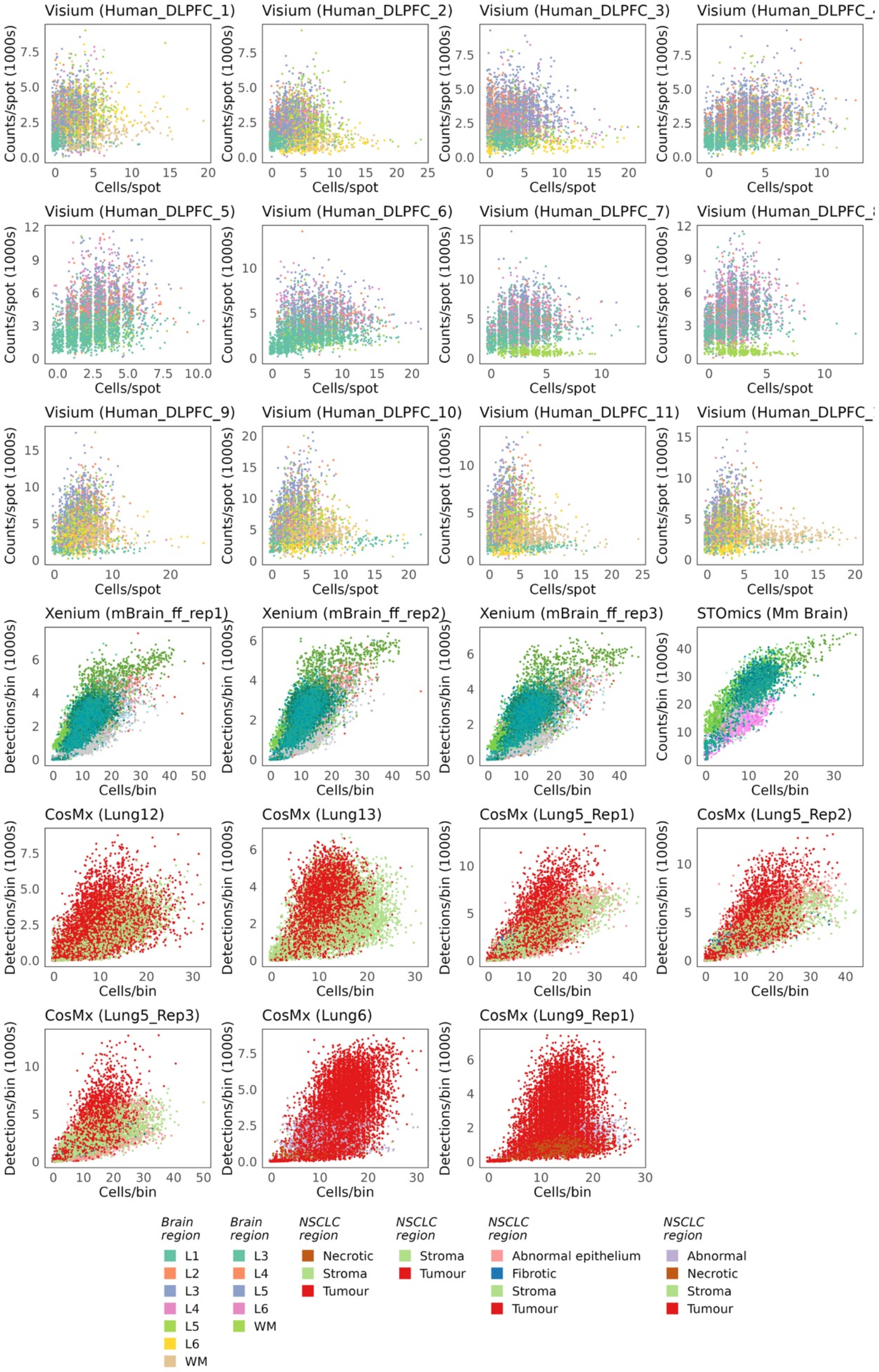


***Supplementary Figure 3:*** *Number of cells plot against the total detections/library sizes per bin/spot, coloured by the tissue region, showing the region-specific relationship between cells and detections/counts.*

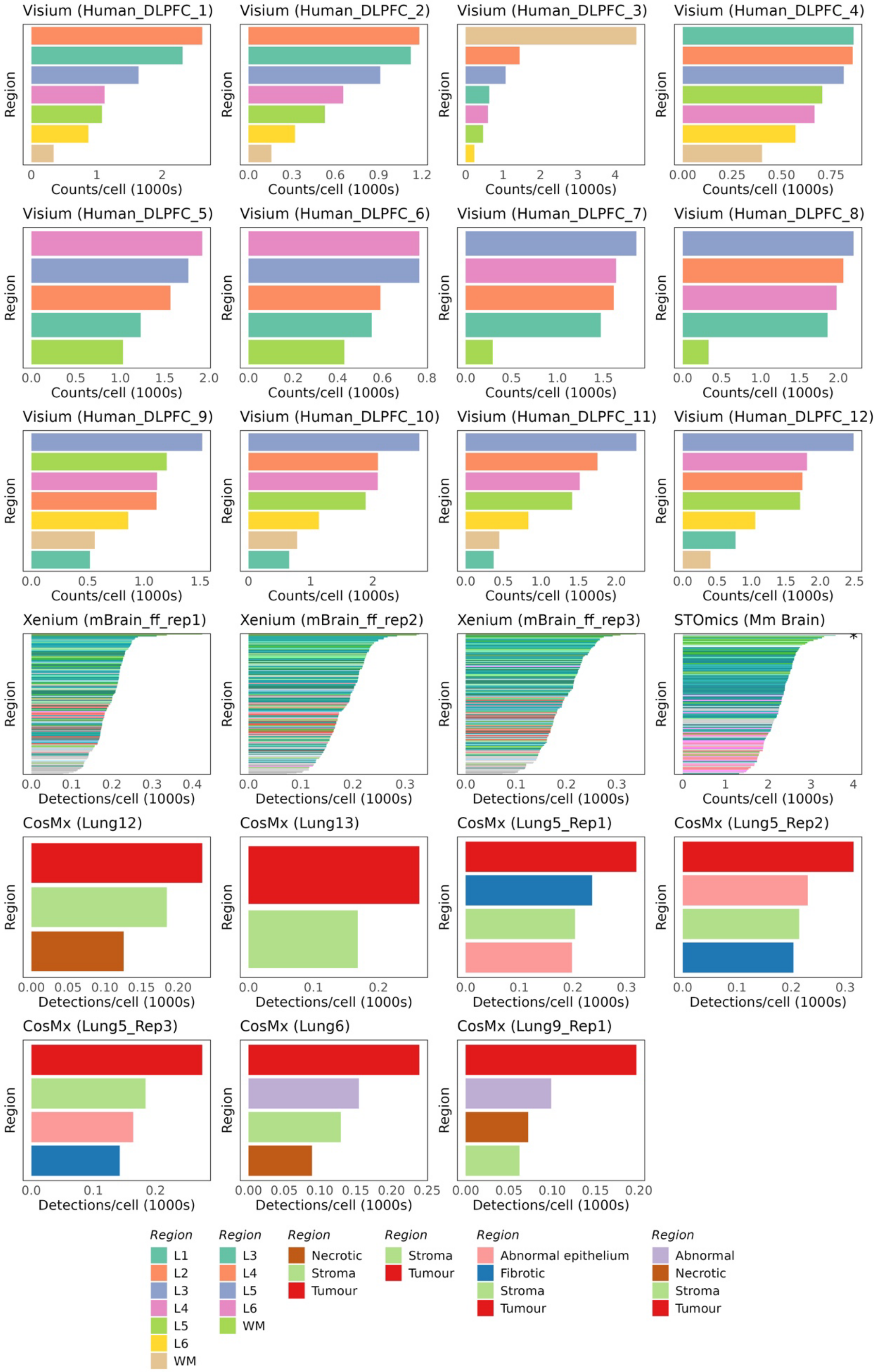


***Supplementary Figure 4:*** *Average detections/library sizes per cell for each region, computed as the sum of detections divided by the number of cells for each region, showing that related regions exhibit similar total detections/library sizes per cell.*
