## Supplementary Table 1 for "Library size confounds biology in spatial transcriptomics data"

| sample | Sum Sq | Df | F value | Pr(>F) | Covariate | Technology |
|---|---|---|---|---|---|---|
| Human_DLPFC_1 | 34500.72457 | 1 | 100.6830114 | 1.98E-23 | NCell | Visium |
| Human_DLPFC_1 | 117179397.4 | 7 | 48851.87095 | 0 | Region | Visium |
| Human_DLPFC_1 | 8845.690978 | 6 | 4.302377302 | 0.000248078 | NCell:Region | Visium |
| Human_DLPFC_2 | 2081.241196 | 1 | 6.837630428 | 0.008956397 | NCell | Visium |
| Human_DLPFC_2 | 92069861.97 | 7 | 43211.8303 | 0 | Region | Visium |
| Human_DLPFC_2 | 26463.78446 | 6 | 14.49051798 | 1.93E-16 | NCell:Region | Visium |
| Human_DLPFC_3 | 7.337372555 | 1 | 0.022607557 | 0.880488452 | NCell | Visium |
| Human_DLPFC_3 | 126789651.4 | 7 | 55808.30509 | 0 | Region | Visium |
| Human_DLPFC_3 | 40029.81576 | 6 | 20.55632173 | 6.75E-24 | NCell:Region | Visium |
| Human_DLPFC_4 | 32441.75496 | 1 | 99.7694206 | 2.96E-23 | NCell | Visium |
| Human_DLPFC_4 | 80591619.22 | 7 | 35406.6593 | 0 | Region | Visium |
| Human_DLPFC_4 | 5147.757496 | 6 | 2.638517262 | 0.01480685 | NCell:Region | Visium |
| Human_DLPFC_5 | 81563.4666 | 1 | 153.1828226 | 1.72E-34 | NCell | Visium |
| Human_DLPFC_5 | 101041666.8 | 5 | 37952.89328 | 0 | Region | Visium |
| Human_DLPFC_5 | 16164.36452 | 4 | 7.589497743 | 4.37E-06 | NCell:Region | Visium |
| Human_DLPFC_6 | 37491.08885 | 1 | 68.34350489 | 1.93E-16 | NCell | Visium |
| Human_DLPFC_6 | 88332967.9 | 5 | 32204.9042 | 0 | Region | Visium |
| Human_DLPFC_6 | 84281.18054 | 4 | 38.40960246 | 1.69E-31 | NCell:Region | Visium |
| Human_DLPFC_7 | 85864.99248 | 1 | 165.9851944 | 2.94E-37 | NCell | Visium |
| Human_DLPFC_7 | 141814970.8 | 5 | 54828.3644 | 0 | Region | Visium |
| Human_DLPFC_7 | 27262.39943 | 4 | 13.17520254 | 1.15E-10 | NCell:Region | Visium |
| Human_DLPFC_8 | 102953.3844 | 1 | 206.0798918 | 1.42E-45 | NCell | Visium |
| Human_DLPFC_8 | 143497872.4 | 5 | 57447.4092 | 0 | Region | Visium |
| Human_DLPFC_8 | 23528.19465 | 4 | 11.77398838 | 1.66E-09 | NCell:Region | Visium |
| Human_DLPFC_9 | 168425.6307 | 1 | 219.4991765 | 2.99E-48 | NCell | Visium |
| Human_DLPFC_9 | 168867149.1 | 7 | 31439.2259 | 0 | Region | Visium |
| Human_DLPFC_9 | 40291.15209 | 6 | 8.751513123 | 1.73E-09 | NCell:Region | Visium |
| Human_DLPFC_10 | 70677.51386 | 1 | 81.42433875 | 2.89E-19 | NCell | Visium |
| Human_DLPFC_10 | 231980298.1 | 7 | 38179.1309 | 0 | Region | Visium |
| Human_DLPFC_10 | 87771.96595 | 6 | 16.85301283 | 2.78E-19 | NCell:Region | Visium |
| Human_DLPFC_11 | 87182.45414 | 1 | 134.5376511 | 1.47E-30 | NCell | Visium |
| Human_DLPFC_11 | 147515245.1 | 7 | 32520.23057 | 0 | Region | Visium |
| Human_DLPFC_11 | 80903.20267 | 6 | 20.80794612 | 4.26E-24 | NCell:Region | Visium |
| Human_DLPFC_12 | 68630.42818 | 1 | 108.4442383 | 5.07E-25 | NCell | Visium |
| Human_DLPFC_12 | 143893562.9 | 7 | 32481.27738 | 0 | Region | Visium |
| Human_DLPFC_12 | 126632.3076 | 6 | 33.3490176 | 2.72E-39 | NCell:Region | Visium |
| mBrain_ff_rep1 | 4898911.503 | 1 | 26560.22025 | 0 | NCell | Xenium |
| mBrain_ff_rep1 | 408444642.2 | 140 | 15817.47825 | 0 | Region | Xenium |
| mBrain_ff_rep1 | 836131.428 | 138 | 32.8494074 | 0 | NCell:Region | Xenium |
| mBrain_ff_rep2 | 5026398.051 | 1 | 28153.66855 | 0 | NCell | Xenium |
| mBrain_ff_rep2 | 377403127 | 144 | 14679.83307 | 0 | Region | Xenium |
| mBrain_ff_rep2 | 1090927.183 | 141 | 43.33659305 | 0 | NCell:Region | Xenium |
| mBrain_ff_rep3 | 4732671.835 | 1 | 24598.76395 | 0 | NCell | Xenium |
| mBrain_ff_rep3 | 372825224.7 | 144 | 13457.04454 | 0 | Region | Xenium |
| mBrain_ff_rep3 | 1044338.793 | 141 | 38.49720427 | 0 | NCell:Region | Xenium |
| STOmics Brain | 5858.958923 | 1 | 136.6888757 | 2.92E-31 | NCell | STOmics |
| STOmics Brain | 64978830.45 | 118 | 12847.02594 | 0 | Region | STOmics |

| | | | | | | |
|---|---|---|---|---|---|---|
| STOmics Brain | 4230921.643 | 1 | 98706.94266 | 0 | NSpots | STOmics |
| STOmics Brain | 38476.17673 | 108 | 8.311528021 | 5.20E-115 | NCell:Region | STOmics |
| STOmics Brain | 62629.93211 | 1 | 1461.149518 | 3.09E-288 | NCell:NSpots | STOmics |
| STOmics Brain | 72415.58257 | 109 | 15.49952 | 1.57E-245 | Region:NSpots | STOmics |
| STOmics Brain | 58914.30558 | 104 | 13.2160034 | 9.20E-197 | NCell:Region:NSpots | STOmics |
| Lung12 | 2259833.315 | 1 | 4861.458487 | 0 | NCell | CosMx |
| Lung12 | 5350074.284 | 3 | 3836.442842 | 0 | Region | CosMx |
| Lung12 | 1632405.46 | 27 | 130.0632043 | 0 | fov | CosMx |
| Lung12 | 30762.27357 | 2 | 33.08861653 | 4.68E-15 | NCell:Region | CosMx |
| Lung12 | 251695.292 | 27 | 20.05402273 | 1.07E-94 | NCell:fov | CosMx |
| Lung12 | 217047.2265 | 26 | 17.9585417 | 1.42E-80 | Region:fov | CosMx |
| Lung12 | 60689.31199 | 26 | 5.021448824 | 8.68E-16 | NCell:Region:fov | CosMx |
| Lung13 | 1014545.21 | 1 | 2156.718714 | 0 | NCell | CosMx |
| Lung13 | 7848359.928 | 2 | 8342.015991 | 0 | Region | CosMx |
| Lung13 | 677814.7917 | 19 | 75.83672034 | 1.19E-276 | fov | CosMx |
| Lung13 | 72.53855549 | 1 | 0.154202354 | 0.694558667 | NCell:Region | CosMx |
| Lung13 | 242284.4564 | 19 | 27.10778635 | 7.45E-95 | NCell:fov | CosMx |
| Lung13 | 169558.6967 | 19 | 18.97092779 | 8.79E-64 | Region:fov | CosMx |
| Lung13 | 52495.97807 | 19 | 5.873466996 | 4.92E-15 | NCell:Region:fov | CosMx |
| Lung5_Rep1 | 5160613.956 | 1 | 10567.5803 | 0 | NCell | CosMx |
| Lung5_Rep1 | 2811192.324 | 4 | 1439.145655 | 0 | Region | CosMx |
| Lung5_Rep1 | 1046522.024 | 29 | 73.89661728 | 0 | fov | CosMx |
| Lung5_Rep1 | 15627.52986 | 3 | 10.66702386 | 5.34E-07 | NCell:Region | CosMx |
| Lung5_Rep1 | 214017.8138 | 29 | 15.11214491 | 4.53E-73 | NCell:fov | CosMx |
| Lung5_Rep1 | 322922.991 | 42 | 15.74431825 | 8.83E-109 | Region:fov | CosMx |
| Lung5_Rep1 | 81919.17259 | 41 | 4.091437152 | 4.17E-17 | NCell:Region:fov | CosMx |
| Lung5_Rep2 | 4728001.116 | 1 | 11554.11108 | 0 | NCell | CosMx |
| Lung5_Rep2 | 2917387.837 | 4 | 1782.350632 | 0 | Region | CosMx |
| Lung5_Rep2 | 941212.4843 | 28 | 82.14641514 | 0 | fov | CosMx |
| Lung5_Rep2 | 12751.27032 | 3 | 10.38702474 | 8.02E-07 | NCell:Region | CosMx |
| Lung5_Rep2 | 133877.4227 | 28 | 11.68445014 | 6.84E-52 | NCell:fov | CosMx |
| Lung5_Rep2 | 168672.2466 | 37 | 11.14040228 | 1.55E-63 | Region:fov | CosMx |
| Lung5_Rep2 | 36668.35533 | 37 | 2.421857996 | 3.16E-06 | NCell:Region:fov | CosMx |
| Lung5_Rep3 | 4346427.84 | 1 | 10273.96546 | 0 | NCell | CosMx |
| Lung5_Rep3 | 2601294.533 | 4 | 1537.21695 | 0 | Region | CosMx |
| Lung5_Rep3 | 1712746.518 | 29 | 139.6049196 | 0 | fov | CosMx |
| Lung5_Rep3 | 24695.56645 | 3 | 19.45823145 | 1.40E-12 | NCell:Region | CosMx |
| Lung5_Rep3 | 150009.0205 | 29 | 12.22714338 | 1.03E-56 | NCell:fov | CosMx |
| Lung5_Rep3 | 197636.2649 | 41 | 11.3943199 | 5.18E-72 | Region:fov | CosMx |
| Lung5_Rep3 | 67589.49816 | 40 | 3.994154482 | 4.19E-16 | NCell:Region:fov | CosMx |
| Lung6 | 1873060.997 | 1 | 4635.317154 | 0 | NCell | CosMx |
| Lung6 | 1558099.719 | 4 | 963.9683875 | 0 | Region | CosMx |
| Lung6 | 4281208.807 | 29 | 365.3389246 | 0 | fov | CosMx |
| Lung6 | 5829.492059 | 3 | 4.808803091 | 0.002387905 | NCell:Region | CosMx |
| Lung6 | 178960.1 | 29 | 15.27164253 | 5.32E-74 | NCell:fov | CosMx |
| Lung6 | 52034.33377 | 17 | 7.574755355 | 4.26E-19 | Region:fov | CosMx |
| Lung6 | 16177.15046 | 16 | 2.502128305 | 0.000781356 | NCell:Region:fov | CosMx |
| Lung9_Rep1 | 949075.956 | 1 | 1453.898786 | 7.31E-300 | NCell | CosMx |

| | | | | | | |
|---|---|---|---|---|---|---|
| Lung9_Rep1 | 4207337.449 | 4 | 1611.315399 | 0 | Region | CosMx |
| Lung9_Rep1 | 1202597.068 | 19 | 96.96159316 | 0 | fov | CosMx |
| Lung9_Rep1 | 2386.661392 | 3 | 1.218716682 | 0.301118668 | NCell:Region | CosMx |
| Lung9_Rep1 | 203611.9555 | 19 | 16.41658716 | 4.51E-54 | NCell:fov | CosMx |
| Lung9_Rep1 | 92510.46577 | 18 | 7.873205103 | 4.19E-21 | Region:fov | CosMx |
| Lung9_Rep1 | 9808.191641 | 15 | 1.001684346 | 0.449692842 | NCell:Region:fov | CosMx |
